# DIFFERENTIAL ANTIPROLIFERATIVE EFFECTS OF CANNABIDIOL (CBD) IN THE CORE AND INFILTRATIVE BOUNDARY OF HUMAN GLIOBLASTOMA CELLS

**DOI:** 10.1101/2024.09.17.613445

**Authors:** Ghazala Abassi-Rana, Yolanda Calle-Patino, Francisco Molina-Holgado

## Abstract

**Background:** We have previously reported that the brain cannabinoid signalling pathways regulates in the isocitrade dehydrogenase-1 wild-type glioblastoma (GBM) core and infiltrative boundary tumor cell proliferation. To uncover the mechanism behind these effects we have investigated the possible antitumoral actions of cannabidiol (CBD) in the tumour core cells (U87) and the Glioma Invasive Margin cells (GIN-8), the latter representing a better proxy of post-surgical residual disease.

**Methods:** Monolayer of GBM cells cultures were treated with increasing concentrations of CBD, Temozolomide (TMZ), Carmustine (BCUN), Fluoxetine, Doxorubicine (DOX) or vehicle. After treatment, cell viability was assessed using an MTT kit assay to evaluate mitochondrial activity/cell proliferation, cytotoxicity was evaluated by LDH release. In addition, we have investigated the effects of the CBD alone or in combination with the above drugs on the autophagic cell death, unfold protein response (UPR) mitochondrial response and release of proinflammatory cytokines.

**Summary:** This study highlights the potential therapeutic relevance of CBD in combination with other FDA-approved drugs against glioblastoma. We observed strong synergism between CBD and TMZ, FX, and DOXO in reducing U87-MG cell viability in vitro, with even stronger synergy between CBD and TMZ in GIN-8 cells. Our preliminary data identify CBD as a potential anti-neoplastic drug in both core and invasive margin cells. Given the heterogeneity of glioblastoma, further studies will elucidate the molecular mechanisms underlying CBD observed anti-tumoral actions and determine whether it can potentially be used in the future as an addition to current therapies.

## Introduction

Brain tumours are the most common tumours that develop in children and adults. Glioblastoma multiforme (GBM) is the most frequent and aggressive primary tumour of the brain, and unlike other tumours, there is currently no cure (Batash et al., 2017), with a survival period post-diagnosis ranging between 12 to 15 months (Ortiz et al., 2019). GBM tumors exhibit resistance to chemotherapy and radiation, with five-year survival rates of approximately four percent (Gomez-Roman et al., 2020). This resistance to conventional treatments has been linked to a subpopulation of cancer cells known as GBM stem-like cells (GSCs) (Gomez-Roman et al., 2020). GSCs demonstrate increased resistance to conventional treatments, including surgical resection, adjuvant radiation, and chemotherapy with temozolomide (TMZ) (Hannan et al., 2019) or lomustine (Doherty et al., 2021).

The tumor microenvironment (TME) offers avenues to identify novel therapeutic targets and strategies to combat GBM or improve the outcome of current treatments (Patterson et al., 2020). The brain endocannabinoid system (ECS) and associated signaling pathways are one such element, which may play a crucial role in shaping the TME in GBM cell lines (Kienzl et al., 2020). Thus, the endocannabinoid system emerges as an important regulator of cell fate, controlling cell survival/cell death decisions depending on the cell type and its environment (Garcia-Arencibia et al., 2019; Hinz and Ramer, 2019).

The ECS comprises endocannabinoids (eCBs), which are endogenous lipid cannabinoids, cannabinoid receptors, and the enzymes responsible for synthesizing and degrading the endocannabinoids (Dasram et al., 2022). Additionally, several phytocannabinoids such as cannabinol (CBN), cannabigerol (CBG), and cannabidiol (CBD) have been studied (Dumitru et al., 2018). CBD is of particular interest for therapeutic purposes related to neurological disorders due to its non-intoxicating properties (Seltzer et al, 2020).

Most cannabinoids bind to the G-protein-coupled cannabinoid receptors CB1 and CB2 (Dumitru et al., 2018). CBD exhibits indirect antagonism of several other G-protein-coupled receptors, including CB1, CB2, and GPR-55 (Thomas et al., 2007; Dumitru et al., 2018). Furthermore, studies have shown that CBD also targets transient receptor potential vanilloid channels TRPV1, TRPV2, and TRPV4 (Santoni and Amantini et al., 2019). The activation of CB1 and CB2 cannabinoid receptors affects various cellular functions (Lu and Mackie, 2016). For instance, cannabinoid receptors inhibit adenylate cyclase, signal through ceramide, and induce mitogen-activated protein kinase (MAPK) and phosphatidylinositol-3-kinase (PI3K) pathways (Dumitru et al., 2018). Both CB1 and CB2 are expressed in glioblastomas (Doherty & De Paula, 2021), with high-grade gliomas, including GBM, exhibiting elevated levels of CB2 (Dumitru et al., 2018).

Certain cannabinoids, such as THC and CBD, have been studied in various glioblastoma models, demonstrating anti-tumoral effects, including the induction of tumor cell death, reduction in tumor cell proliferation, and modulation of angiogenesis (Doherty & De Paula, 2021; Feng et al., 2024). Despite these anti-tumoral effects, one of the only FDA-approved drugs containing CBD for GBM in clinical trials is Sativex, which contains THC and CBD in a 1:1 ratio (Doherty et al., 2021). Other FDA-approved drugs for GBM, such as Doxorubicin and Carmustine, have been employed for treatment, albeit with limited efficacy.

In this study, FDA-approved drugs for GBM, including temozolomide (TMZ), fluoxetine (FX), carmustine (BCNU), and doxorubicin (DOXO), were co-incubated with CBD in 2D GBM in vitro models to analyze potential synergy between the drugs and their anti-tumoral effects, aiming to arrest tumor progression. Additionally, we explored the potential signaling mechanisms through which CBD targets the tumorigenicity of GBM. GBM cells from the core and invasive margins respond differently to radiotherapy. Several studies have shown that U87-MG core cells are intrinsically more radioresistant and less susceptible to radiotherapy compared to the more invasive U251 cells, which exhibit enhanced migratory potential (He et al., 2022). Unfortunately, GBM also presents heterogeneous responses to drug therapy, with drug penetration through the blood-brain barrier (BBB) being a dose-limiting factor (Vasey et al., 2021). Therefore, post-surgical intracranial drug delivery strategies, such as polymer pro-drug nanoparticles (NP), are being developed to ensure localized drug delivery within the brain (Vasey et al., 2021).

Interestingly, cell lines that represent the more residual cancerous cells left in the brain after maximal safe surgical resection, such as GIN-8, GIN-28, and GIN-31, show higher NP uptake compared to the U87-MG classical core tumor cell line (Vasey et al., 2021). Furthermore, GIN-28 cells were found to be the most susceptible to DOX-based treatments, followed by GIN-8 and GIN-31, which exhibited similar drug sensitivity (Vasey et al., 2021). However, U87-MG cells were the most treatment-sensitive cell line under both free DOX and DOX-NP conditions (Vasey et al., 2021). This may be due to the lower proliferative rate of patient-derived primary cell lines compared to the in vitro-adapted U87-MG cell line (Vasey et al., 2021). Consequently, the GIN invasive margin lines represent a more accurate model for testing therapeutic response in GBM, especially given that around 85 percent of GBMs relapse locally within 2 cm³ of the infiltrative margin post-standard combination therapy with oral TMZ and radiotherapy (Vasey et al., 2021).

To uncover the biological mechanisms behind CBD’s anti-tumor activity, we conducted a detailed investigation into the effects of FDA-approved drug combinations with CBD on brain cancer cell fate. This included a thorough analysis of differences between U87-MG core and GIN-8 invasive margin cell lines. Specifically, we performed assays to evaluate the neuroimmune scenario (expression and function of inflammatory cytokines), involvement of autophagy, and the role of the mitochondrial unfolded protein response (UPRmt) in CBD-treated U87-MG and GIN-8 cell lines.

## Methods

### Reagents

(-)-Cannabidiol (CBD) was purchased from Tocris (Bristol, UK). All other reagents and materials for cell cultures were obtained from standard suppliers.

### Cellular Models

All glioblastoma cell lines (U87 and GIN-8) used in this project were of human origin and obtained from Dr. Ruman Rahman (University of Nottingham). U87 cells, confirmed by Short Tandem Repeat (STR) genotyping, were isolated from the core of a GBM tumor and sourced commercially, serving as a biological positive control for GBM cells. The Glioma Invasive Margin (GIN-8) cell line, isolated from the tumor’s infiltrative edge, was derived in-house from surgeries at the Queen’s Medical Centre, Nottingham (and confirmed to be comparable to their respective primary tissue by STR analysis).

Monolayer cell cultures were prepared as described previously (Williams et al, 2022). Briefly, cells were plated into T75 cell culture flasks (Nunc, UK) until they reached confluency. Cells were then trypsinized and plated at a density of 25,000 cells/ml (Figure 1) on different plastic platforms in DMEM (Sigma, UK), supplemented with 10% fetal bovine serum (Sigma, UK), 5mM sodium pyruvate (Sigma, UK), and 5mM L-glutamine (Sigma, UK). The cultures were maintained in a humidified incubator at 37°C and 5% CO₂. Cells grown in 24-well, 48-well, or 96-well plates were treated with increasing concentrations of CBD, temozolomide (TMZ), carmustine (BCNU), fluoxetine, doxorubicin (DOX), or vehicle.

**Figure 1.**
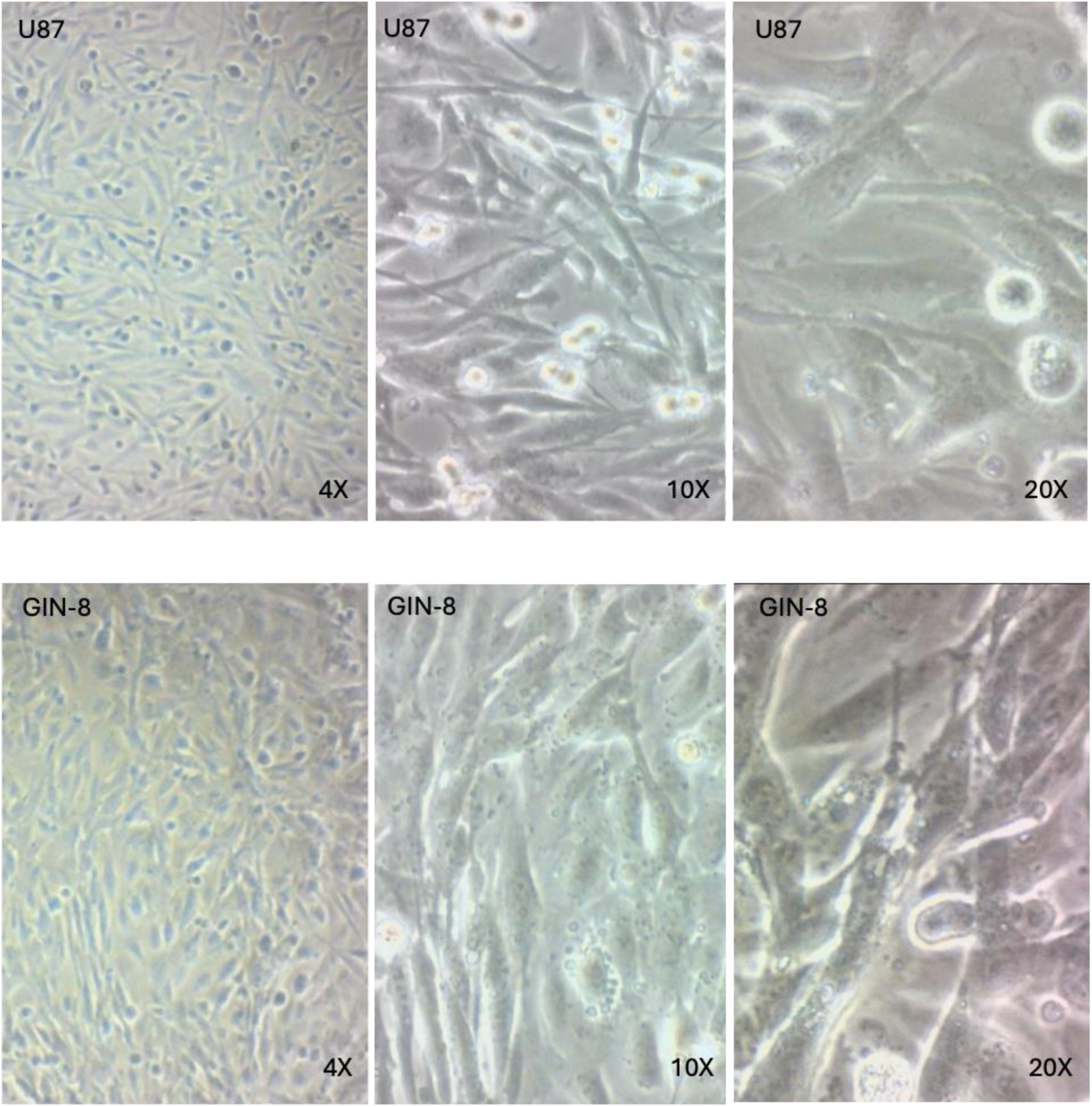
Representative images of morphology of human glioblastoma cells. The upper panel shows phase contrast images of untreated monocultured U87 cells at different magnifications (4x, 10x, and 20x). The lower panel presents phase contrast micrographs of untreated monocultured GIN-8 cells.

Cytotoxicity was evaluated by measuring the release of the cytosolic enzyme LDH into the culture medium, a marker of dead and dying cells, using the CytoTox-96 LDH assay (Promega, UK). Cell viability was assessed with an MTT kit assay (Sigma-Aldrich, UK) to evaluate the production of formazan blue, which indicates mitochondrial activity and cell proliferation. Additionally, we evaluated the effects of CBD alone or in combination with the above drugs on cell proliferation, cytotoxicity, autophagic cell death, unfolded protein response (UPR), mitochondrial response, and the release of various proinflammatory cytokines (IL-1β, TNF-α, IFN-β, IFN-γ, and IL-6).

### Assessment of cellular viability (MTT)

Cell viability was assessed using an MTT assay which is a marker of mitochondrial activity (Carmichael et al. 1987). Following exposure of neuronal cultures to CBD or different antitumoral drugs, the cultures were washed twice with sterile phosphate-buffered saline before the addition of MTT (1 mg/mL) in HBM pH 7.4 (5 mM HEPES, 154 mM NaCl, 4.6 mM KCl, 2.3 mM CaCl2, 33 mM glucose, 5 mM NaHCO3, 1.1 mM MgCl2 and 1.2 mM Na2HPO4). Following incubation (45 min; 37°C), MTT solutions were removed, and the formazan product was solubilized in DMSO, and the absorbance read at 505 nm using a Versemax plate reader. Data were presented as a percentage relative to their vehicle controls. All data from MTT measurements are expressed as Formazan blue production % of control, considered as 100%.

### Assessment of cell death (LDH)

Cytotoxicity was evaluated by release of the cytosolic enzyme LDH into the culture medium by dead and dying cells (CytoTox-96 LDH assay; Promega, Southampton, UK). Total LDH release was calculated by incubating untreated cells with 0.1% Triton X-100 for 10 min (37°C, 5% CO2, 95% air) to induce maximal cell lysis. Absorbance was measured at 490 nm using a Versemax plate reader. Treatment values were then expressed as a percentage of the total LDH release. Background LDH release (media alone) was subtracted from the experimental values. All data from LDH measurements are expressed as % cytotoxicity, where 100% = maximum toxicity induced by the insult.

### Autophagy Assay

The autophagic activity in cell cultures and its response to different drug treatments was assessed using an autophagy assay kit (Sigma, Missouri, USA) following the manufacturer’s instructions. In brief, 20μL of autophagosome detection reagent was diluted in 10ml stain buffer in a 96 well plate. Once the medium was removed from controls and samples, 100μL of autophagosome detection reagent working solution was added. The cells were then incubated for 1hr. Following incubation, the cells were washed with 100μL of wash buffer, and plates were then read at 520nm using a microplate reader.

### ELISA Assays

TNF-α was measured using a validated, specific sandwich enzyme-linked immunosorbent assay (ELISA; Campbell et al., 1997) purchased from R&D Systems (UK). IL-1β was measured by ELISA using kits purchased from R&D Systems (UK). The assays were specific for human TNF-α or human IL-1β and there was no cross reactivity with other cytokines. The sensitivity of the TNF-α kit was 4 pg/ml whereas that of the IL-1β kit was 1 pg/ml.

### Nuclear extraction

Nuclear extracts were prepared with the EpiSeeker Nuclear Extraction Kit (Signosis, Inc.) according to the user manual. The cells were washed twice in phosphate-buffered saline (PBS) and lysed on ice for 10 min in the extraction buffer I with gentle shaking, and then were collected from the plates, and centrifuged at 15,000 rpm for 3 min at 4 °C. The supernatant (cytoplasmic fraction) was discarded; the pellet was then resuspended in 250 μl of extraction buffer II and incubated on ice for 2 h with gentle shaking. After the mixture was centrifuged at 15,000 rpm for 5 min at 4 °C, the supernatant containing nuclear protein was collected and ready for assays.

### Mitochondrial UPR transcription factors (TF) Activation Profiling Array

The mitochondrial unfolded protein response (UPRmt) in CBD-treated U87-MG and GIN-8 cell lines] was assessed using profiling plate array kits to detect mitochondrial UPR transcription factors (TF) activation (Signosis, Inc., USA). Each array assay was performed according to the procedure described in the Mitochondrial UPR TF Activation Profiling Plate Array kit user manual. In brief, 10 μg of nuclear extract was first incubated with the biotin-labeled probe mix at room temperature for 30 minutes. The activated TFs bound to their corresponding DNA binding probes. After isolating the protein/DNA complexes from unbound probes, the bound probes were eluted and hybridized with the plate pre-coated with capture oligonucleotides. The captured biotin-labeled probes were then detected with streptavidin–HRP and subsequently measured using a chemiluminescent plate reader (Veritas microplate luminometer).

## RESULTS

### Anti-proliferative Effects of Cannabidiol (CBD) in the Core and Invasive Margin of Human Glioblastoma Cells

To assess the differential anti-proliferative effects of cannabidiol (CBD) on the core and invasive margin of human glioblastoma (GBM) cells, we conducted a series of cell viability and proliferation assays on CBD-treated GBM cell cultures (U87 and GIN-8). In both cell lines, CBD effectively reduced the viability of GBM cells. CBD treatment resulted in a significant, concentration-dependent reduction in viability for both U87 cells (Figure 2A) and the more clinically relevant invasive-margin GIN-8 cells (Figure 2B).

**Figure 2.**
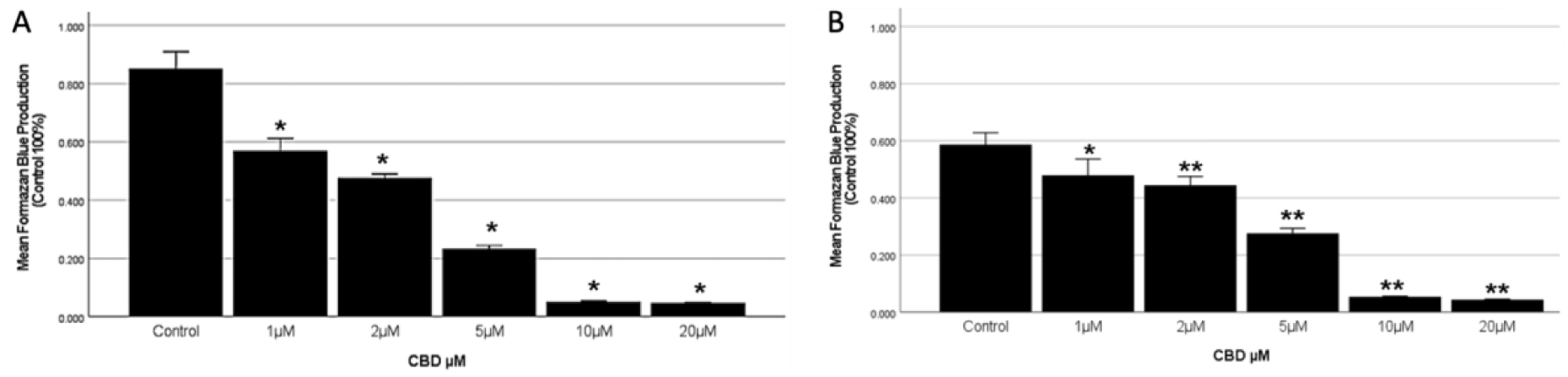
**(A)** Effect of CBD on the proliferation of U87 core glioblastoma (GBM) cells. The potential cytotoxic effect of CBD was evaluated in adult human U87 glioblastoma cells seeded at a density of 25,000 cells/ml and exposed to different CBD concentrations (1µM, 2µM, 5µM, 10µM and 20uM) or vehicle (t=24h) through a MTT assay. Statistics: \**p* < 0.01 versus control. The results shown are the mean ± SEM of six independent experiments in quadruplicate. **(B)** Effect of CBD on the proliferation of the GIN-8 GBM invasive margin cells. The cytotoxic effect of CBD was evaluated in GIN-8 cells (25,000 cells/ml) and exposed to different CBD concentrations (1µM, 2µM, 5µM, 10µM and 20uM) or vehicle (t=24h) through a MTT assay. Statistics: \**p* < 0.005; ** *p* < 0.001 *versus* control. The results shown are the mean ± SEM of six independent experiments in quadruplicate.

### Cannabidiol (CBD) induces cell death in the Core and Invasive Margin of Human Glioblastoma Cells

To assess the differential cytotoxic effects of cannabidiol (CBD) on the core and invasive margin of human glioblastoma (GBM) cells, we evaluated the release of the cytosolic enzyme LDH into the culture medium by dead and dying cells in CBD-treated GBM cell cultures (U87 and GIN-8). In both cell lines, CBD effectively increased cytotoxicity, reducing the viability of GBM cells. CBD treatment resulted in a significant, concentration-dependent increase in cytotoxicity for both U87 cells (Figure 3A) and the invasive-margin GIN-8 cells (Figure 3B).

**Figure 3.**
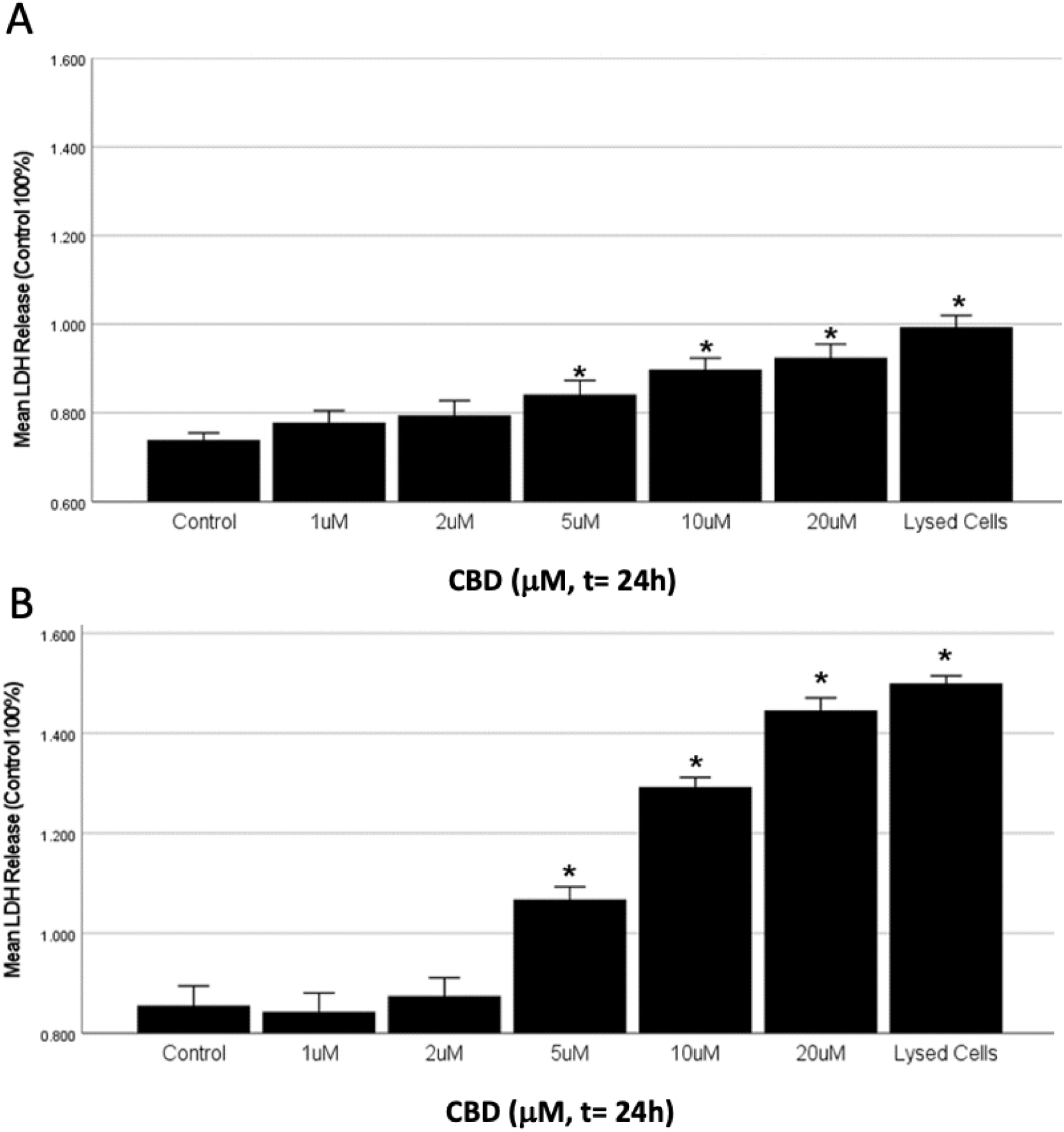
**(A)** Cytotoxicity of CBD on the proliferation of U87 core glioblastoma (GBM) cells was evaluated by release of the cytosolic enzyme LDH into the culture medium by dead and dying cells (CytoTox-96 LDH assay; Promega, Southampton, UK). U87 glioblastoma cells were seeded at a density of 25,000 cells/ml and exposed to different CBD concentrations or vehicle. Statistics: \**p* < 0.001 versus control. The results shown are the mean ± SEM of six independent experiments in quadruplicate. Total LDH release was calculated by incubating untreated cells with 0.1% Triton X-100 for 10 min (37°C, 5% CO 2, 95% air) to induce maximal cell lysis. **(B)** Cytotoxicity of CBD on the proliferation of the GIN-8 GBM invasive margin cells. The cytotoxic effect of CBD was evaluated in GIN-8 cells (25,000 cells/ml; t=24h) and exposed to different CBD concentrations or vehicl through a LDH assay. Statistics: \**p* < 0.001 *versus* control. The results shown are the mean ± SEM of six independent experiments in quadruplicate.

### CBD induced autophagic cell death in human glioblastoma cells

Following our observation that CBD reduces the viability of both U87-MG and GIN-8 cells, we investigated how CBD affects these cells and aimed to identify the underlying cellular mechanisms. We specifically evaluated the role of CBD-induced lethal autophagy in U87-MG (Figure 4A) and GIN-8 cells (Figure 4B). Our results demonstrated that CBD treatment led to a dose-dependent increase in autophagy (*p < 0.001), peaking at 5 µM CBD and slightly declining at concentrations of 10 µM and higher in both U87-MG and GIN-8 cells. However, autophagy levels were significantly higher in U87-MG cells at 10 µM and 20 µM CBD compared to the invasive-margin GIN-8 cells. The observed decrease in autophagy at 10 µM CBD may be due to a high percentage of cells undergoing CBD-induced cell death, leaving only a small fraction detectable by the autophagy assay.

**Figure 4.**
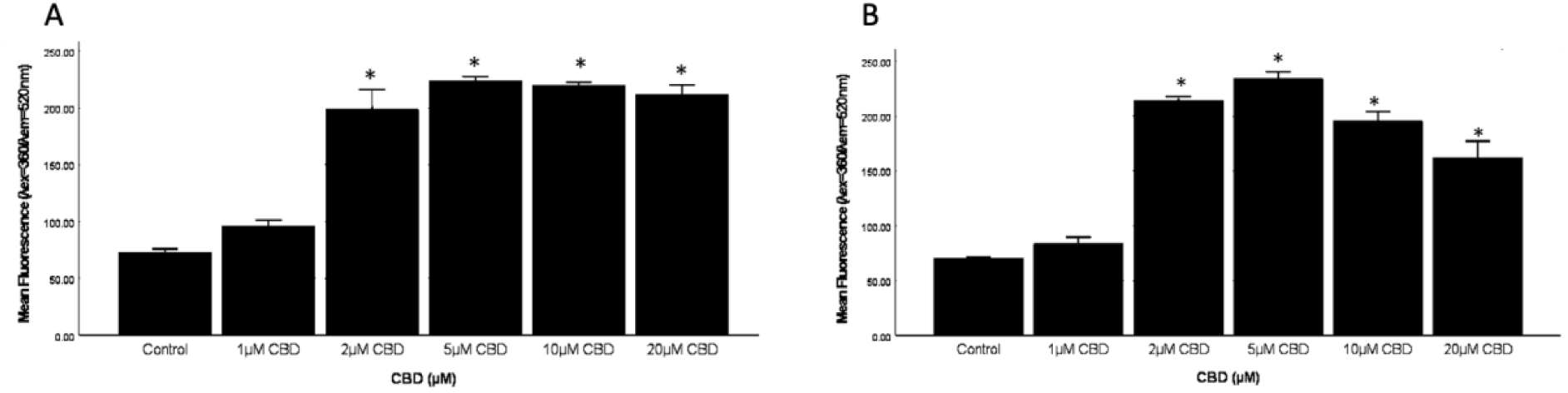
Autophagy was measured in U87 cells, plated at 25000 cells/ml in a 96 well plate, by an autophagy assay using a propriety fluorescent autophagosome marker (λex = 333/λem = 518 nm) after 24 hrs of CBD treatment. **(A)** Evaluation of autophagic cell death in CBD treated U87-MG cells. Statistics: *p < 0.001 versus control. The results shown are the mean ± SEM of six independent experiments in quadruplicate. (B) CBD-induced autophagic cell death in GIN-8 cells. Statistics: *p < 0.001 versus control. The results shown are the mean ± SEM of six independent experiments in quadruplicate.

### Differential effects of CBD exposure on the mitochondrial unfolded protein response (UPR-mt) in the Core and Invasive Margin of Human Glioblastoma Cells

We investigated the involvement of the mitochondrial unfolded protein response (UPR-mt) in the pathophysiology of GBM by analyzing the activation of various related transcription factors (TFs) in CBD-treated U87-MG core and invasive margin GIN-8 cells (Figures 5 and 6) using a mitochondrial activation profiling plate array. The experiments involved plate array analysis of 16 TFs in U87 and GIN-8 cells, treated with or without CBD. After 24 hours of treatment, cells were harvested, and nuclear extracts were analyzed. Our findings revealed significant variability in the activation of UPR-mt-associated transcription factors across U87-MG core and invasive margin GIN-8 cell samples. For example, basal levels of transcription factor activation appeared higher in GIN-8 cells compared to U87-MG cells. However, when assessing the change in TF activation between untreated and CBD-treated samples for each cell line, we observed that the difference in TF activation was significantly greater in U87-MG cells in response to CBD.

**Figure 5.**
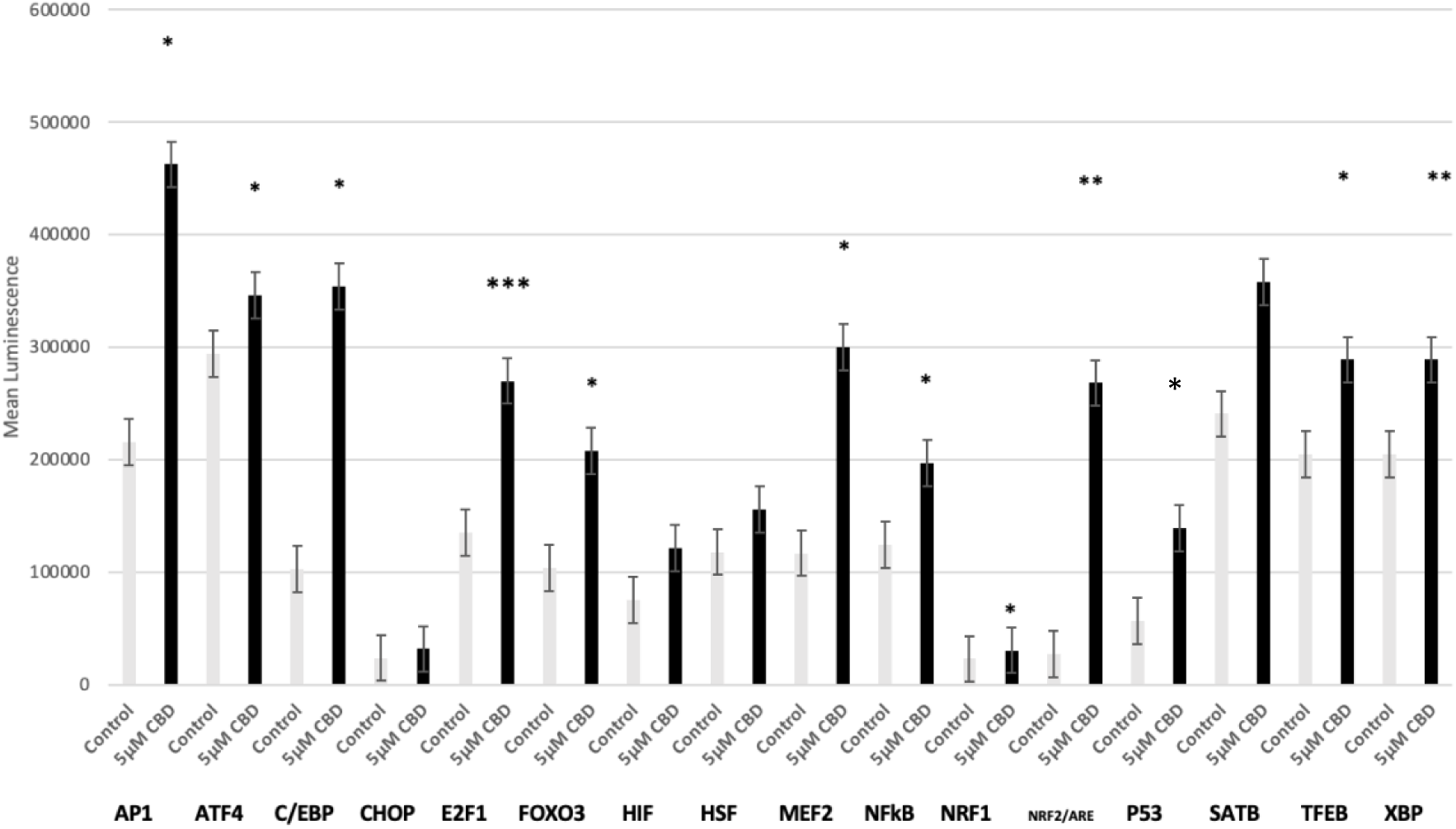
Activation of UPR-mt related Transcription Factors in CBD treated U-87 Cells. The mitochondrial UPR Transcription Factors were monitored in nuclear extracts of U87-MG cells treated with CBD (5µM, t=24h, 25,000 cells/mL). Statistics: *p ≤ 0.05 vs Control; **p ≤ 0.01 vs. Control; ***p ≤ 0.001 vs Control. The results shown are the mean ± SEM of four independent experiments in triplicate.

**Figure 6.**
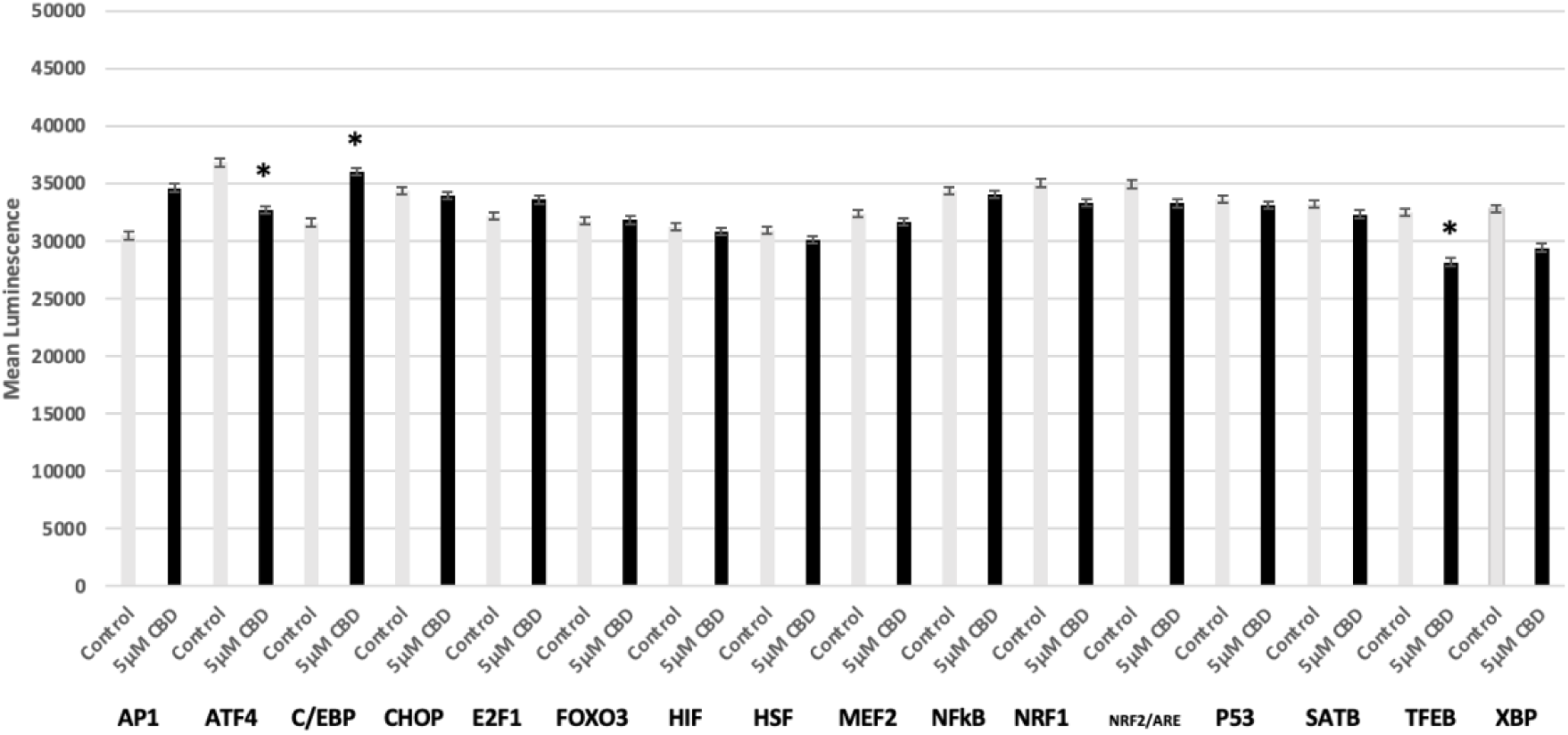
Activation of the mitochondrial unfold protein response (UPR-mt) related transcription factors (TF) in CBD treated GIN-8 Cells. UPR-mt TF were monitored in nuclear extracts of GIN-8 cells treated with CBD (5µM, t=24h, 25,000 cells/mL). Statistics: *p ≤ 0.05 vs Control. The results shown are the mean ± SEM of four independent experiments in triplicate.

Additionally, we found a marked increase in p53 activation in CBD-treated U87-MG cells compared to GIN-8 cells. Other transcription factors activated by CBD in U87-MG but not in GIN-8 cells included nuclear factor kappa B (NF-κB) and Special AT-rich sequence-binding protein 2 (SATB2). In contrast, Activating Transcription Factor 4 (ATF4) was significantly expressed in both CBD-treated U87-MG and GIN-8 cells. The differential activation of transcription factors between U87-MG core and invasive margin GIN-8 cells suggests that CBD may utilize distinct pathways to target these two GBM cell lines.

### CBD enhances Temozolomide (TMZ) cytotoxic effect in U87-MG and GIN-8 cells

Another series of MTT assays was conducted on U87-MG and GIN-8 cells, where the cell cultures were co-incubated with CBD and TMZ to evaluate the effects of both drugs on the viability of core and invasive margin GBM cells, both independently and in combination. The results demonstrated a pronounced cytotoxic effect of TMZ on U87-MG (Figure 7A) and GIN-8 cells (Figure 7B), which intensified when co-incubated with CBD in a dose-dependent manner. Moreover, combinations of TMZ and CBD produced moderate to strong antagonistic effects in the invasive margin GIN-8 cells, whereas they resulted in strong synergistic antitumoral effects in the core U87-MG cells. These effects were observed for both lower and higher doses of TMZ, with examples including 125 µM TMZ and 5 µM CBD or 500 µM TMZ and 10 µM CBD in vitro.

**Figure 7.**
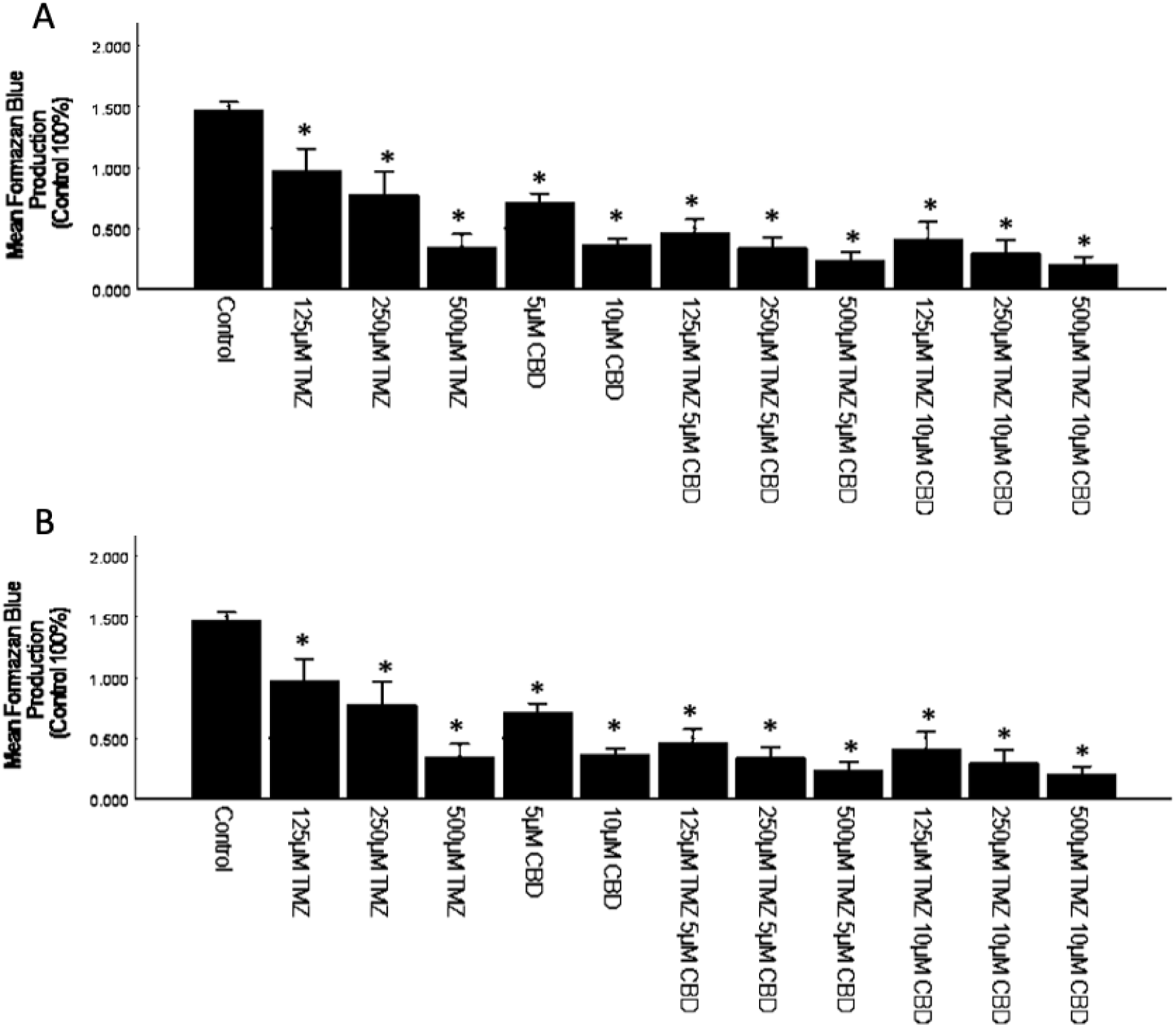
**(A)** Effect of CBD and TMZ on the proliferation of U87-MG cells. The cytotoxic effects of TMZ alone co-incubated with CBD were evaluated in adult human U87-MG cells (25000 cells/ml; t=24h) through an MTT analysis. Statistics: * p < 0.001 *vs* control; # p < 0.001 cells co-treated with TMZ and CBD vs cells treated with TMZ. The results shown are the mean ± SEM of six independent experiments in quadruplicate. (B) Effect of CBD and TMZ on the proliferation of GIN-8 cells (25000 cells/ml; t=24h; n=6 in quadruplicate). Statistics: * p < 0.001 *vs* control.

### CBD enhances Fluoxetine (FX) cytotoxic effect in U87-MG and GIN-8 cells

The MTT assay was repeated to test the effects of CBD and fluoxetine (FX) on the viability of core U87-MG (Figure 8A) and invasive margin GIN-8 cell cultures (Figure 8B). The effects of each drug were tested independently as well as in combination. Both drugs lowered the viability of each cell line in a concentration-dependent manner, whether used alone or together. CBD significantly intensified the cytotoxic effect of FX on the viability of both cell lines, with the U87-MG core cell line being slightly more sensitive to the drugs compared to the invasive margin GIN-8 cells. A significant decrease in U87-MG cell viability was observed with low doses of 5 µM CBD and 10 µM FX in combination (Figure 8A). Interestingly, combinations of fluoxetine (FX) and CBD demonstrated synergistic antitumoral effects in both cell lines. Specifically, synergistic effects were observed at doses of 15 µM FX and 5 µM CBD in GIN-8 cells, and 20 µM FX and 10 µM CBD in U87 cells.

**Figure 8.**
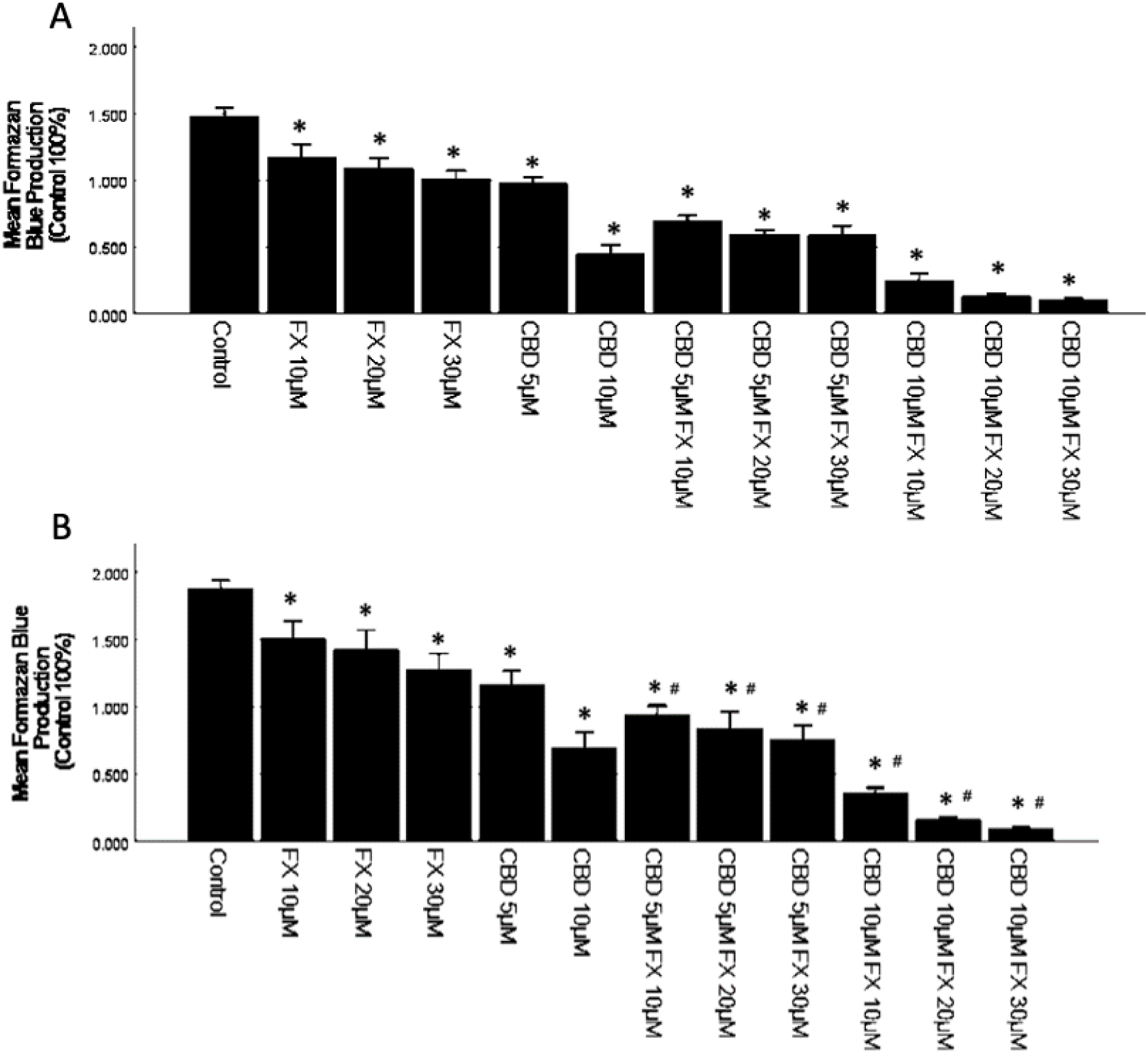
**(A)** Effect of CBD and FX on the proliferation of U87-MG cells. The cytotoxic effects of FX alone co-incubated with CBD were evaluated in adult human U87-MG cells (25000 cells/ml; t=24h) through an MTT analysis. Statistics:* p < 0.001 *vs* control; # p < 0.001 cells co-treated with FX and CBD vs cells treated with FX. The results shown are the mean ± SEM of six independent experiments in quadruplicate. **(B)** Effect of CBD and FX on the proliferation of GIN-8 cells (25000 cells/ml; t=24h; n=6 in quadruplicate). Statistics: *p < 0.001 *vs* control; # p < 0.001 cells co-treated with FX and CBD vs cells treated with FX.

### CBD enhances Carmustine (BCUN) cytotoxic effect in U87-MG and GIN-8 cells

Co-incubations of CBD and BCNU were also tested to evaluate their effects on the cell viability of both core U87-MG cells (Figure 9A) and invasive GIN-8 cells (Figure 9B). The results showed BCNU to be one of the most cytotoxic drugs tested in this study, causing a significant decrease in cell viability in both U87-MG and GIN-8 cells, whether used alone or in combination with CBD. The combination of CBD (5 µM) with BCNU (100 µM) resulted in a synergistic antitumoral effect in the invasive margin GIN-8 cells but not in the core U87-MG cells.

**Figure 9.**
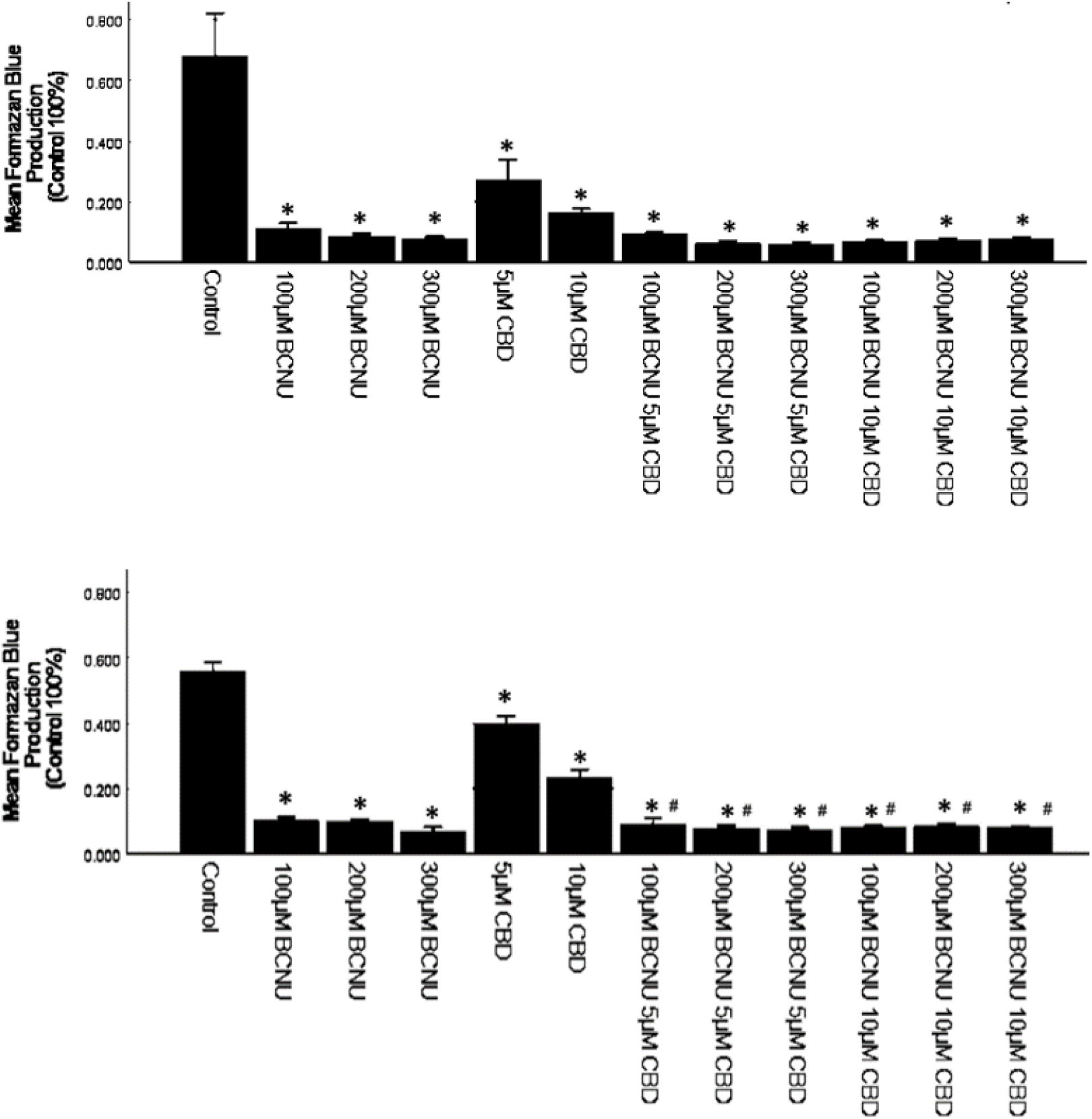
**(A)** Effect of CBD and Carmustine (BCNU) on the proliferation of U87-MG cells. The cytotoxic effects of BCNU alone co-incubated with CBD were evaluated in adult human U87-MG cells (25000 cells/ml; t=24h) through an MTT analysis. Statistics: *p < 0.001 *vs* control; # p < 0.001 cells co-treated with BCUN and CBD vs cells treated with BCUN. The results shown are the mean ± SEM of six independent experiments in quadruplicate. **(B)** Effect of CBD and BCNU on the proliferation of GIN-8 cells (25000 cells/ml; t=24h; n=6 in quadruplicate). Statistics: *p < 0.001 *vs* control; # p < 0.001 cells co-treated with BCNU and CBD vs cells treated with BCNU.

### CBD enhances Doxorubicin (DOXO) cytotoxic effect in U87-MG and GIN-8 cells

Proliferation assays on both core U87-MG and invasive margin GIN-8 cell cultures were conducted to study the effects of CBD and DOXO, both independently and in co-incubation (Figure 10). Consistent with the results obtained with other drugs tested in this study, both CBD and DOXO caused a concentration-dependent decrease in the viability of each cell line. While incubation with DOXO alone led to a significant reduction in cell viability, it did not result in a substantial decrease. However, co-incubation of CBD and DOXO in both cell cultures resulted in a more pronounced reduction in the viability of GBM cells. Specifically, co-treatment with 10 µM CBD and 5 µM Doxorubicin (DOXO) in vitro resulted in a synergistic antitumoral effect in both the core U87-MG and invasive margin GIN-8 cells.

**Figure 10.**
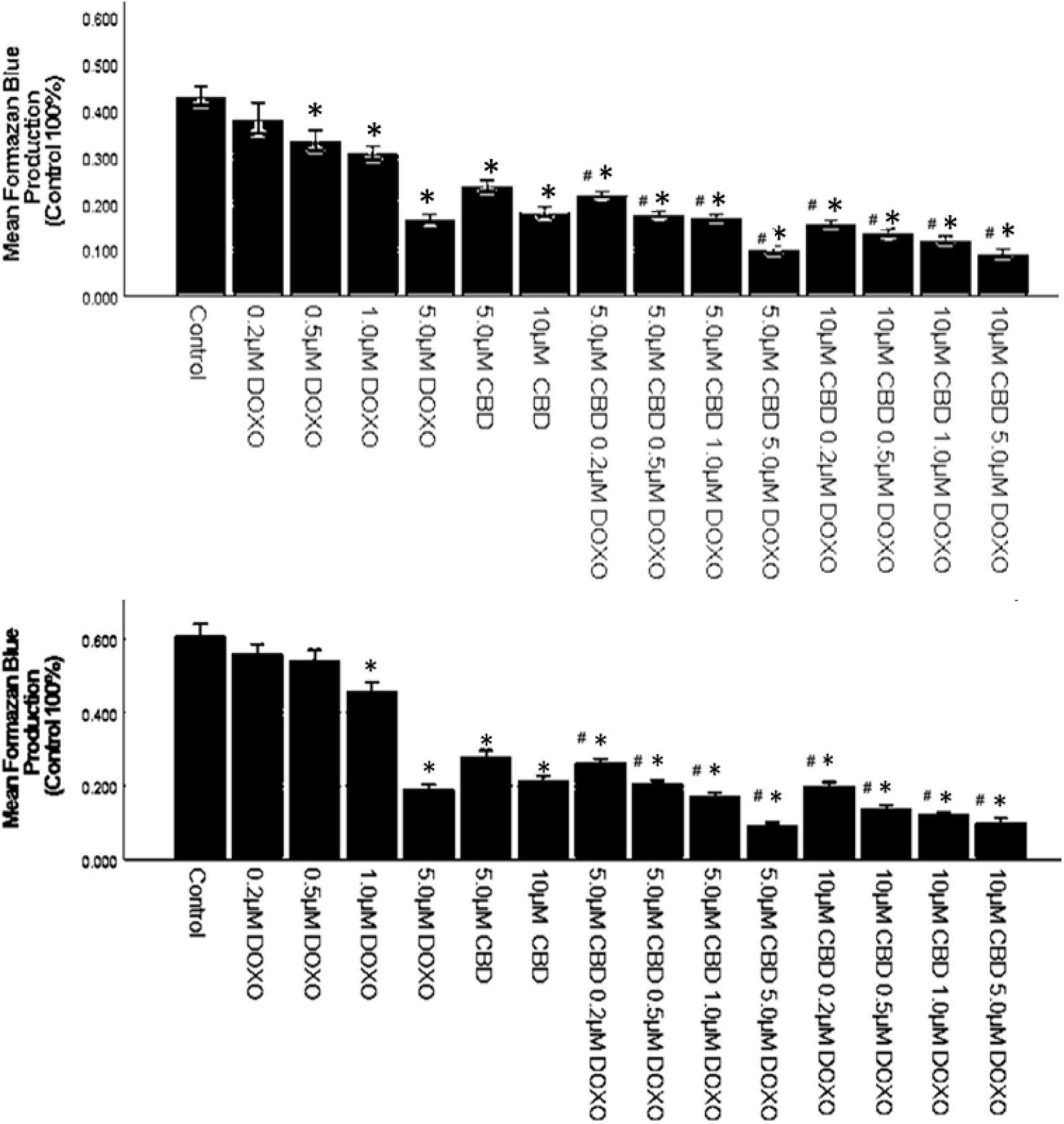
**(A)** Effect of CBD and Doxoribucin (DOXO) on the proliferation of U87-MG cells. The cytotoxic effects of BCNU alone co-incubated with CBD were evaluated in adult human U87-MG cells (25000 cells/ml; t=24h) through an MTT analysis. Statistics: *p < 0.001 *vs* control; # p < 0.001 cells co-treated with DOXO and CBD vs cells treated with DOXO. The results shown are the mean ± SEM of six independent experiments in quadruplicate. **(B)** Effect of CBD and DOXO on the proliferation of GIN-8 cells (25000 cells/ml; t=24h; n=6 in quadruplicate). Statistics: *p < 0.001 *vs* control; # p < 0.001 cells co-treated with DOXO and CBD vs cells treated with DOXO.

### Differential Neuroimmune Effects of CBD on Cytokine Production in Core and Invasive Margin Human Glioblastoma Cells

To understand the neuroimmune scenario associated with CBD-induced cell death in both core U87-MG and invasive margin GIN-8 cells, a series of ELISA assays were performed to investigate cytokine secretion in CBD-treated U87-MG and GIN-8 cells (Figures 11 and 12). Both core and invasive margin GBM cells were treated with CBD at varying concentrations for 24 hours. ELISA assays were then conducted to evaluate the secretion of pro-inflammatory cytokines such as TNF-α, IL-1β, IFN-β, IL-6, and IFN-γ in CBD-treated U87-MG and GIN-8 cells.

**Figure 11.**
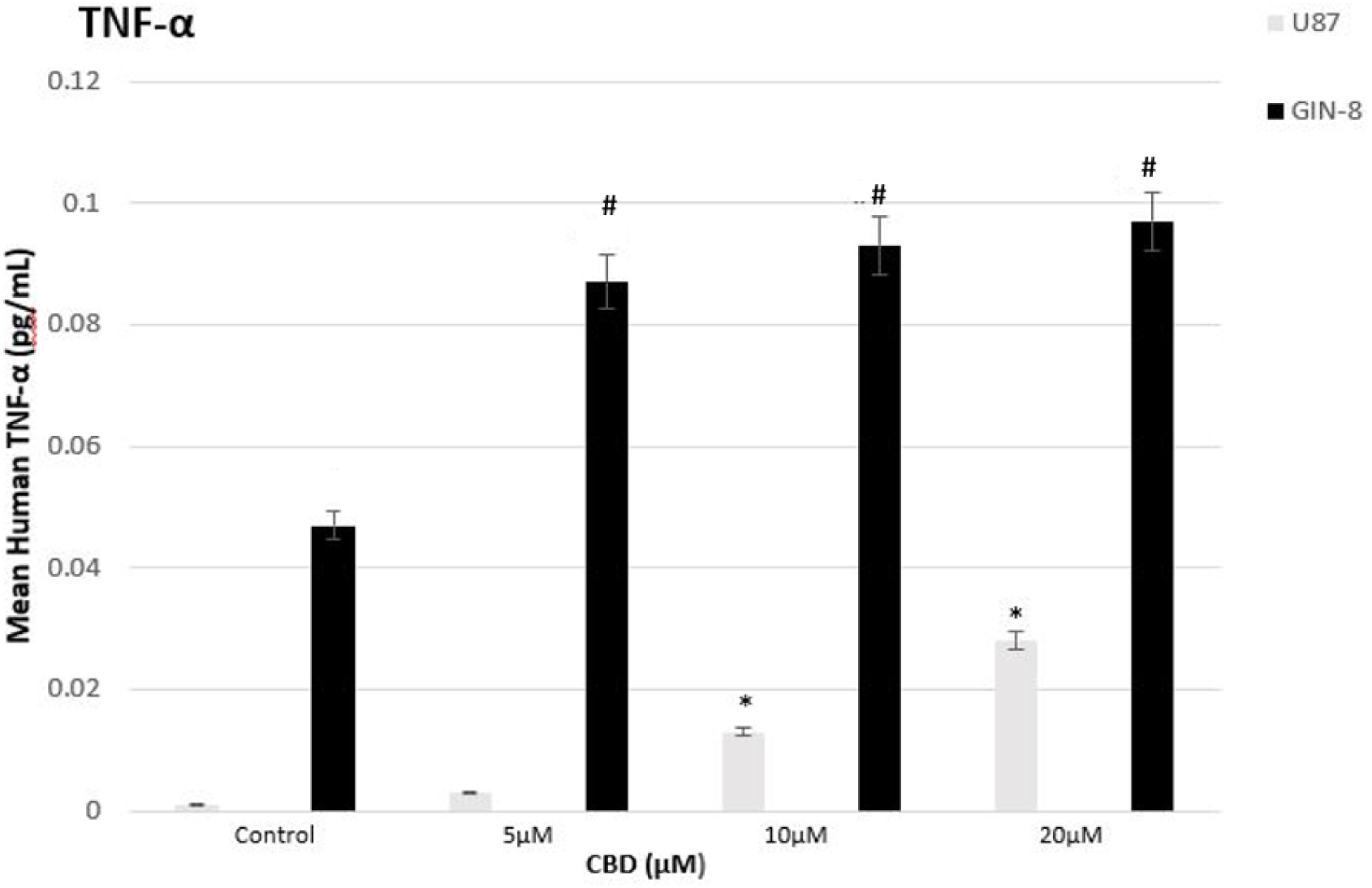
CBD upregulates TNF-α production in GBM cells. U-87 and GIN-8 cell cultures (25000 cells/ml) were treated for 24 h with CBD, supernatants collected and the levels of TNF-α were measured by ELISA. Data are presented as mean±SEM of six independent determinations. Statistical differences: U87: ^*^;p < 0.001 vs. Control; GIN-8: ^#^p < 0.001 vs Control.

**Figure 12.**
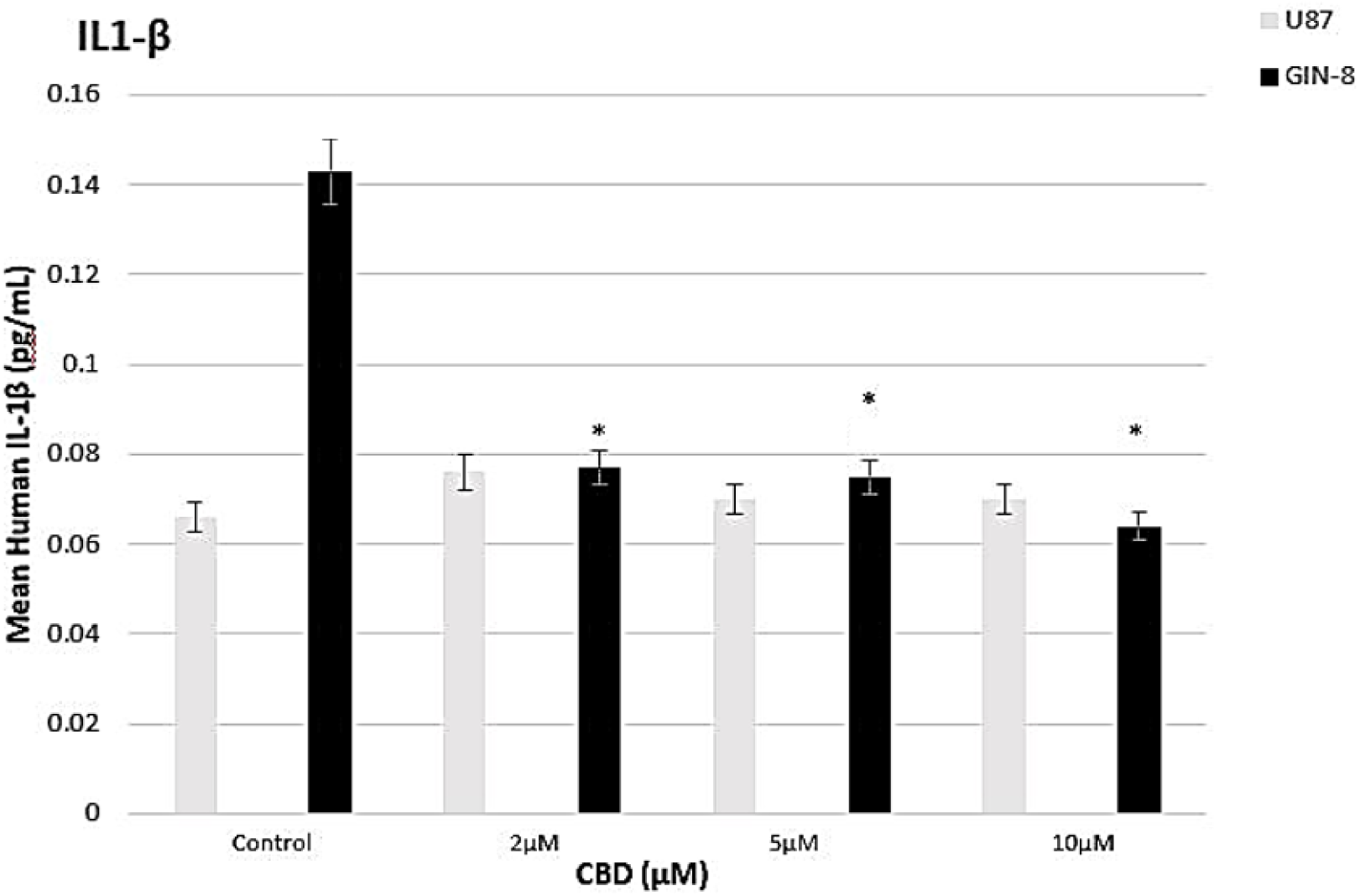
Effects of CBD on the IL-1β production in GBM cells. U-87 and GIN-8 cell cultures (25000 cells/ml) were treated for 24 h with CBD, supernatants collected and the levels of IL-1β were measured by ELISA. CBD down-regulates the production of IL-1β in U87 cells, Data are presented as mean±SEM of six independent determinations. Statistical differences: U87: ^*^p < 0.001 vs. Control;

We observed that CBD triggered a significant, concentration-dependent increase in TNF-α secretion in both U87-MG and GIN-8 cells. Notably, significantly higher levels of TNF-α were observed in all CBD-treated GIN-8 cell groups compared to the U87-MG cell groups. In contrast, CBD induced a significant, concentration-dependent decrease in IL-1β secretion in GIN-8 cells. These results suggest that CBD modulates pro-inflammatory and pro-tumoral signaling pathways through IL-1β in GBM cells (Tarassishin et al., 2014). GIN-8 cells exhibited significantly higher basal levels of IL-1β compared to U87-MG cells. CBD had no effect on the basal levels of IFN-γ, IL-6, and IFN-β in either U87-MG or GIN-8 cells (data not shown).

## DISCUSSION

Different signaling pathways associated with the brain’s endocannabinoid system are being investigated as potential therapeutic target (Velasco et al., 2012). Recent evidence suggests that blocking the cannabinoid system mediates antitumor effects via the inactivation of traditional cannabinoid receptors (CB1 and CB2) (Malfitano et al., 2012; Proto et al, 2012). However, the molecular mechanisms through which CBD affects glioblastoma (GBM) cells remain unclear and require further investigation. Given the heterogeneity exhibited by GBM cells, it is essential to identify the properties of not only core tumor cells but also the more aggressive and invasive margin cells to develop therapeutic strategies suited to each cell type. This field is still developing, with limited results, as the molecular mechanisms of the antitumor effects downstream of cannabinoid receptor activation are not yet fully understood (Fraguas et al., 2018).

In this context, we previously reported the effectiveness of the CB2 receptor inverse agonist AM630 in blocking the proliferation of core GBM cells (Williams et al., 2022). However, AM630 appears to be a less potent inhibitor of the invasive margin cell population.

The main findings of this study suggest that CBD differentially inhibits the proliferation of both core and invasive glioblastoma cells in vitro. One potential mechanism involved is the activation of autophagy by CBD in both core and invasive glioblastoma cells. This aligns with and expands on previous studies demonstrating that CBD-driven autophagy is mediated by the CB1, CB2, TRPV1(Vrechi et al., 2021), and GPR55 receptor (Lah et al., 2022). Our study confirms and extends these findings, as we observed iCB1, CB2, and GPR55 receptor expression in both core and invasive glioblastoma cells (data not shown).

Our results also revealed that CBD-induced autophagic cell death occurs in a concentration-dependent manner. For instance, autophagy peaked at 5 μM CBD in vitro for both the U87-MG core and GIN-8 invasive glioblastoma cell lines. Several studies have shown a correlation between CBD treatment and increased autophagy in GBM cells via ERK activation and reactive oxigeen species production (Kim et al., 2024). For example, one study observed higher levels of the autophagy marker LC3A in glioblastoma cells treated with higher CBD concentrations (Soroceanu et al., 2022). In our study, most U87-MG and GIN-8 cells die in culture when treated with CBD concentrations above 10μM, which may explain the significant decrease in autophagy in both cell lines at 10μM and 20μM CBD. Nonetheless, several research studies indicate that autophagy plays a dual role in GBM cells, acting as both protective and lethal depending on the stress conditions (Yun et al., 2021). Glioblastoma cells exhibit higher basal autophagy levels, which help them survive the early stages of stress, but autophagy decreases once cellular resources are depleted (Ivanov et al., 2020). While we investigated the increased sensitivity to CBD-induced autophagy in core and invasive margin GBM cells in this study, future research could focus on whether this autophagy upregulation is protective or lethal by suppressing autophagic flux in core and invasive GBM cultures.

Moreover, studies have shown that CBD induces mitochondrial dysfunction and lethal mitophagy arrest, leading to autophagic glioblastoma cell death (Huang et al., 2021). For instance, it was found that CBD reduces mitochondrial quality and quantity in glioblastoma cells, inducing PINK1 translocation to the mitochondrial membrane, triggering mitophagy. These results indicated that endoplasmic reticulum (ER) stress is a key regulator of CBD-induced mitophagy in glioblastoma cells (Huang et al., 2021).

To further understand the role of mitochondrial dysfunction and stress response in CBD-induced autophagic cell death, we performed a mitochondrial unfolded protein response (UPRmt) assay on CBD-treated U87-MG core and GIN-8 invasive cells. Mitochondrial activity is highly elevated in glioblastoma and other cancerous cells due to uncontrolled proliferation and nutrient shortages (Altea-Manzano et al., 2020). Several studies have demonstrated the cytotoxic nature of CBD in GBM cells, suggesting that CBD dysregulates mitochondrial activity (Gross et al., 2021). This dysregulation causes significant stress, activating the UPRmt to maintain protein homeostasis by upregulating chaperones and enhancing the mitochondria’s folding capacity during stress (Riess et al., 2022). Based on the cytotoxic nature of CBD observed in our U87-MG core and GIN-8 invasive cell lines, we hypothesized that the UPRmt pathway is also involved in these cells’ response to CBD. Our study identified several transcription factors activated by UPRmt in both CBD-treated U87-MG core and GIN-8 invasive cells, with more factors activated in U87-MG cells compared to GIN-8 cells. For example, p53 was significantly activated in CBD-treated U87-MG cells but not in GIN-8 cells, suggesting heterogeneity in GBM cell responses to CBD. It is possible that CBD induces p53 pathway modifications in U87-MG cells, potentially activating p53-dependent apoptotic pathways in GBM cells with wild-type p53(Ivanov et al., 2017). However, studies have also found a correlation between p53 mutations and GBM progression (Hosami et al., 2021), disrupting mitochondrial apoptotic pathways (Ivanov et al., 2017). These findings highlight the variability in how GBM cell lines respond to CBD treatment.

In addition to p53, the nuclear factor kappa B (NF-kB) was activated in CBD-treated U87-MG cells. Previous research has shown that NF-kB promotes tumor progression in GBM cells by regulating metabolic adaptation and sustaining cell viability. Interestingly, recent studies have shown that CBD is able to convert NF-κB into a tumor suppressor in glioblastoma (Volmar et al., 2021). This may support the clinical implementation of CBD as a promising drug for GBM and the development of new NF-kB-modulating compounds for cancer therapy.

The transcription factor SATB2 was also highly expressed in CBD-treated U87-MG cells. SATB2 is enriched in GBM and promotes tumor growth by enhancing GSC proliferation through the recruitment of histone acetyltransferase CBP, leading to increased FOXM1 expression (Tao et al., 2020).

Another transcription factor, ATF4, was significantly expressed in both CBD-treated U87-MG and GIN-8 cells. Previous studies have identified the ATF4-DDIT3-TRIB3-Akt-mTOR axis as involved in CBD-induced mitophagy in GBM cells (Huang et al., 2021). Our results support the hypothesis that autophagic cell death occurs in both core and invasive GBM cells treated with CBD.

Pro-inflammatory cytokines also play a key role in GBM pathophysiology (Propper et al., 2022). Moreover, cytokines can alter the anti-tumor response in the context of brain metastases, suggesting that cytokine networks can be manipulated for diagnosis and treatment. (Fares et al. 2021). Our findings show that TNF-α is significantly secreted in both CBD-treated U87-MG and GIN-8 cells, with GIN-8 cells exhibiting higher basal TNF-α levels. Our findings are in agreement and further expand previous studies that identified the cellular tumor antigen p53, NF-kappa-B, and TNF-α as lead targets o CBD activity (Ma et al., 2021).

Additionally, CBD-treated GIN-8 cells secreted high levels of IL-1β at basal levels, which declined in a concentration-dependent manner upon CBD treatment. This suggests that CBD activates the JNK signaling pathway to induce autophagy as a mechanism of cell death in both core and invasive GBM cells (Ivanov et al., 2017).

In breast cancer cells, CBD is able to revert the mesenchymal invasive phenotype of cancr cells induced by the Inflammatory IL-1β (Garcia-Morales et al, 2020). This ability of CBD may be highly useful in GBM therapy, as tumor cells treated with CBD become more sensitive to conventional brain cancer treatments. Based on our findings, this could be important for reducing cell viability and inhibiting the migration of invasive margin GBM cells.

Lastly, our study highlights the potential therapeutic relevance of CBD in combination with other FDA-approved drugs against glioblastoma. We observed a strong synergism between CBD and TMZ, FX, and DOXO in reducing U87-MG cell viability in vitro, with even stronger synergy between CBD and TMZ in GIN-8 cells. In support of the above, in experimental animal models this combined therapy of CBD and TMZ has shown to exert antitumoral actions in several types of cancer including glioma (Velasco et al., 2018).

This suggests that combining CBD with existing treatments could be clinically significant in combating GBM and the present findings may inform such developments. In conclusion, this study provides novel insights into the mechanistic actions and pathways through which CBD combats GBM by comparing its effects on core and invasive glioblastoma cells. These findings lay the groundwork for future studies to further evaluate CBD effects on the GBM tumor microenvironment and explore potential therapeutic targets in more complex 2D and 3D culture systems. Given the heterogeneity of glioblastoma, more studies are required to elucidate the molecular mechanisms underlying CBD observed anti-tumoral actions and determine whether it can potentially be used in the future as an addition to current therapies.

## Authors Contribution

All Authors have made a substantial contribution to this project and have approved the submitted version of the manuscript.

